# Repression of MUC1 promotes expansion and suppressive function of myeloid-derived suppressor cells in pancreatic ductal adenocarcinoma and breast cancer murine models

**DOI:** 10.1101/2020.12.07.415299

**Authors:** S. Mahnaz, L. Das Roy, M. Bose, C. De, S. Nath, P. Mukherjee

## Abstract

Myeloid-derived suppressor cells (MDSCs) are immature myeloid cells that are responsible for immunosuppression in tumor microenvironment. Here we report the impact of mucin 1 (MUC1), a transmembrane glycoprotein, on proliferation and functional activity of MDSCs. To determine the role of MUC1 in MDSC phenotype, we analyzed MDSCs derived from wild type (WT) and MUC1-knockout (MUC1KO) mice bearing pancreatic ductal adenocarcinoma KCKO and breast cancer C57MG xenografts. We observed enhanced tumor growth in MUC1KO mice compared to WT mice in both pancreatic KCKO and breast C57MG cancer models due to increased MDSC population and enrichment of Tregs in tumor microenvironment. Our current study shows that knockdown of MUC1 in MDSCs promotes proliferation and immature suppressive phenotype indicated by increased level of iNOS, ARG1 activity and TGF-β secretion under cancer conditions. Increased activity of MDSCs leads to repression of IL-2 and IFN-ɣ production by T-cells. We were able to find that MDSCs from MUC1KO mice have higher levels of c-Myc and activated pSTAT3 as compared to MUC1 WT mice, that are signaling pathways leading to increased survival, proliferation and prevention of maturation. In summary, MUC1 regulates signaling pathways that maintain immunosuppressive properties of MDSCs. Thus, immunotherapy must target only tumor associated MUC1 on epithelial cells and not MUC1 on hematopoietic cells to avoid expansion and suppressive functions of MDSC.

## INTRODUCTION

A diverse population of immature myeloid cells (IMCs) make up the Myeloid-derived suppressor cells (MDSCs). This heterogenous population consists of cells (IMCs) with precursors for macrophages, granulocytes or dendritic cells (DCs) that accumulate in chronic inflammation and tumor progression [1–4]. In mice, MDSCs express both CD11b and Gr1 markers and mainly are of two subtypes: polymorphonuclear Ly6G^+^Ly6C^lo^ (PMN) and monocytic Ly6G^−^Ly6C^hi^ (M) cells. In humans, these same subtypes can be characterized as Lin^−^HLA-DR^−^/^lo^CD33+ or Lin-HLA-DR-/^lo^CD11b^+^CD14^−^CD15^+^CD33^+^ for PMN-MDSCs and CD14^+^HLA^−^DR^neg^/^lo^ or Lin^−^HLA^−^DR^neg^/^lo^CD11b^+^CD14^+^CD15^−^ for M-MDSCs [1, 2, 5–7]. MDSCs are derived from the hematopoietic precursor cells in the bone marrow through modulation of myelopoiesis by inflammatory mediators [1–3, 5, 8]. They exhibit highly immunosuppressive and tumorigenic activities [1–3, 9]. MDSCs suppress anti-tumor immune response by diverse mechanisms, including deprivation of amino acids arginine and cysteine, essential for T-cell proliferation and anti-tumor immune response [1, 10, 11], production of nitric oxide (NO) and reactive oxygen species (ROS) leading to nitration of T-cell receptors (TCR) and chemokines necessary for T-cell migration and induction of apoptosis of T cells and NK cells [1–3, 12, 13]; production of interleukin (IL)-10 and transforming growth factor β 1 (TGFβ-1) that inhibit functions of immune effector cells [1–3, 10, 14], increasing expression of programmed death-ligand 1 (PD-L1) [1–3, 15] which can downregulate anti-tumor T cell-functions by interacting with PD-1 receptor expressed on T cells [16], secretion of angiogenic factors like VEGF [17, 18], and growth factors, matrix metalloproteinases and cytokines promoting tumor growth and activation of regulatory T cells (Tregs) [2, 19, 20]. Therefore, MDSCs are important players in tumor-mediated immunosuppression. MDSCs fail to differentiate into macrophages, granulocytes and dendritic cells under cancer conditions [21, 22]. In murine tumor models, co-expression of the CD11b and myeloid cell lineage differentiation antigen GR1 is a distinctive MDSC phenotype marker [23]. MDSCs were reported as a major obstacle in achieving response to immune therapy [24].

Many studies have demonstrated that after the MDSCs migrate to the tumor site, there is a significant upregulation in their immunosuppressive functions. Inflammatory cytokines like interferon-γ (IFN-γ), interleukins 1, 4 and 13 (IL-1, IL-4, IL-13), tumor-necrosis factor-α (TNF-α), toll-like receptor (TLR) ligand and prostaglandin E2 (PGE2) provide these activating signals that are mediated by STAT1, STAT6, NF-κB and by an increased activity of cyclooxygenase 2(COX-2) [1, 2, 10, 25–28]. COX2 has been shown to be regulated by mucin protein MUC1 in pancreatic cancer [29].

Another study showed that development of bone marrow (BM) progenitors into CD11b+Gr1+ MDSCs is dependent on down-regulation of β–catenin levels, which is also regulated by MUC1 [30]. MUC1 is a transmembrane glycoprotein that is expressed on the glandular or luminal epithelial cells of the mammary gland, esophagus, stomach, duodenum, pancreas, uterus, prostate, and lungs, and in hematopoietic cells [31, 32]. In normal cells, MUC1 is only expressed on the apical surface and is heavily glycosylated with the core protein sequestered by the carbohydrates. As cells start transforming to a malignant phenotype, expression of MUC1 increases by many folds, and the expression is no longer restricted to the apical surface, but it is found all around the cell surface and in the cytoplasm [33]. In addition, glycosylation on tumor-associated MUC1 (tMUC1) is aberrant, with greater exposure of the peptide core than is found in normal tissues [34]. MUC1 was ranked as the second most optimal target out of 75 for immunotherapy by the National Cancer Institute [35]. Previously we have shown that MUC1 expression in the tumor helps in maintaining the MDSCs in an immature and immunosuppressive state, leading to an aggressive nature of MUC1+ PDA tumors [36]. This makes tMUC1 on epithelial tumors an emerging target for immunotherapy [37, 38]. However, non-specific targeting leads to side-effects. Although MUC1 is known to be an epithelial cell marker, its expression and function on normal and neoplastic hematopoietic cells have been described [39, 40]. The role of MUC1 in immature myeloid cells should be explored to further understand its regulatory mechanisms and also find ways to avoid undesired consequences of targeting MUC1 on normal MDSCs.

In this study, we investigated the expansion, differentiation, and suppressive function of MDSCs in MUC1-wild type (WT) and MUC1-null mice (MUC1KO) under cancer conditions. Using two models of epithelial originated cancers, namely, pancreatic ductal adenocarcinoma (PDA) KCKO and breast cancer C57MG cells, we have shown that not only MUC1KO mice have higher levels of MDSC expansion in the spleen, but also that these MDSCs have more suppressive phenotype compared to that of MDSCs in WT mice. Since systemic targeting of MUC1 in many carcinomas is being actively explored [41], the fact that MUC1 is involved in a regulatory mechanism of expansion and immunosuppressive activity of MDSCs is of great importance. From our studies, it becomes clear that therapeutic targeting of MUC1 must be addressed specifically to the epithelial tumors because downregulation of MUC1 expression in hematopoietic cells will lead to increased MDSCs with immunosuppressive phenotype.

## MATERIALS AND METHODS

### Cell lines

PDA cell line KCKO was generated from spontaneous PDA MUC1-null tumors and maintained as previously described [42]. C57MG breast cancer cells were kindly provided by Dr. Gendler, Mayo clinic and maintained according to the published protocol [43].

### Mouse model

Male C57BL/6 mice were obtained from The Jackson Laboratory and MUC1KO mice were generated as previously described [44]. Mice were handled and maintained in accordance with the University of North Carolina at Charlotte Institutional Animal Care and Use Committee approved protocol. Mice were 3 months old at the time of tumor injection. For the PDA model, 10^6^ KCKO cells were injected in the right flank of the mouse. For the breast cancer model, 10^6^ C57MG cells were injected in the right low mammary fat pad. Palpable tumors were measured by calipers every two days, and tumor weight was calculated according to the following formula: milligrams = (length × (width)2)/2, where length and width are measured in centimeters.

### Flow cytometry

5×10^5^ bone marrow cells were obtained from the femurs and tibias of mice. Splenocytes from control and tumor bearing mice were derived as previously described [36]. Tumor infiltrating lymphocytes were isolated by digesting tumors with 0.5mg/ml Collagenase III (Worthingthon) for 20 min at 37°C. Tumors were incubated with DNAse I for 10 min at room temperature. Samples were passed through a 40uM screen and finally centrifuged in 40/60 Percoll for 20 min. Lymphocytes were collected and then stained with fluorescence-conjugated antibodies. Acquisition was performed by BD FACS Calibur and BD Fortessa. Data analysis was done using FLOWJo (Ashland, OR).

### Antibodies

Fluorescent antibodies against Gr1 (Miltenyi Biotec), CD11b (BD), iNOS (BD), Arginase-1, c-Myc (SantaCruz Biotech), Sca1 (Stem Cell Technologies), MUC1 (BD), pSTAT3 (eBioscience) and isotype control (eBioscience) were used at a concentration of 1μg/sample.

### Arginase assay

2×10^5^ MDSCs were subjected to Arginase activity assay. Briefly, cells were lysed with 1% Triton-X in 37°C. 25mM Tris-HCl and 10mM MnCl_2_ were added to samples. Lysates were then heated at 56°C to activate Arginase-1. 0.5M L-arginine was added to the lysates and the reaction was stopped using phosphoric and sulphuric acid. 40 μl of a-isonitrosopropiophenone was added and samples were heated at 95°C for 30 minutes. Samples were read at 450 nm.

### ELISA

Levels of cytokine TGF-β in the murine serum was analyzed with a kit from eBioscience using sandwich ELISA.

### MDSC suppression assay

CD3^+^T-cells were isolated from healthy C57/B6 mice using a Miltenyi kit (Miltenyi Biotec, Auburn, CA). Round bottom plates were coated with 10μg of functional CD3 antibody (eBiosciences, San Diego, CA) and 5 μg CD28 antibody overnight. 10^5^ CD3+T-cells were plated in RPMI containing 10% heat inactivated fetal calf serum, 1% penicillin/streptomycin and 1% glutamax (Invitrogen). MDSCs were sorted first by enriching the CD11b+ cells from the spleen and then sorted for Gr1+cells using a Miltenyi kit. 10^5^ MDSCs were incubated with CD3+T-cells for 72 hours. Supernatants were collected at 72 hours and were then subjected to IL-2 and IFN-ɣ ELISA assays (eBiosciences).

### Statistical analysis

Statistical analysis was performed with GraphPad Prism software. One-way ANOVA was performed between the groups and the significance was confirmed using the Duncan and Student-Newman-Keul test. Values were considered significant if p < 0.05.

## RESULTS

### MUC1KO mice are more susceptible to tumor growth

MDSCs are an important component of tumor immunosuppressive microenvironment causing susceptibility to tumor growth. In our study, a comparison of tumor growth and final wet weight between WT and MUC1KO mice indicate that in the global absence of MUC1, mice are more susceptible to tumor growth with both KCKO and C57MG cell lines. In fact, not only tumors grow at a significantly faster rate in MUC1KO mice (Figure 1A, C), but also tumors from WT mice never match the weight of that of MUC1KO mice (Figure 1B, D).

**Fig 1.**
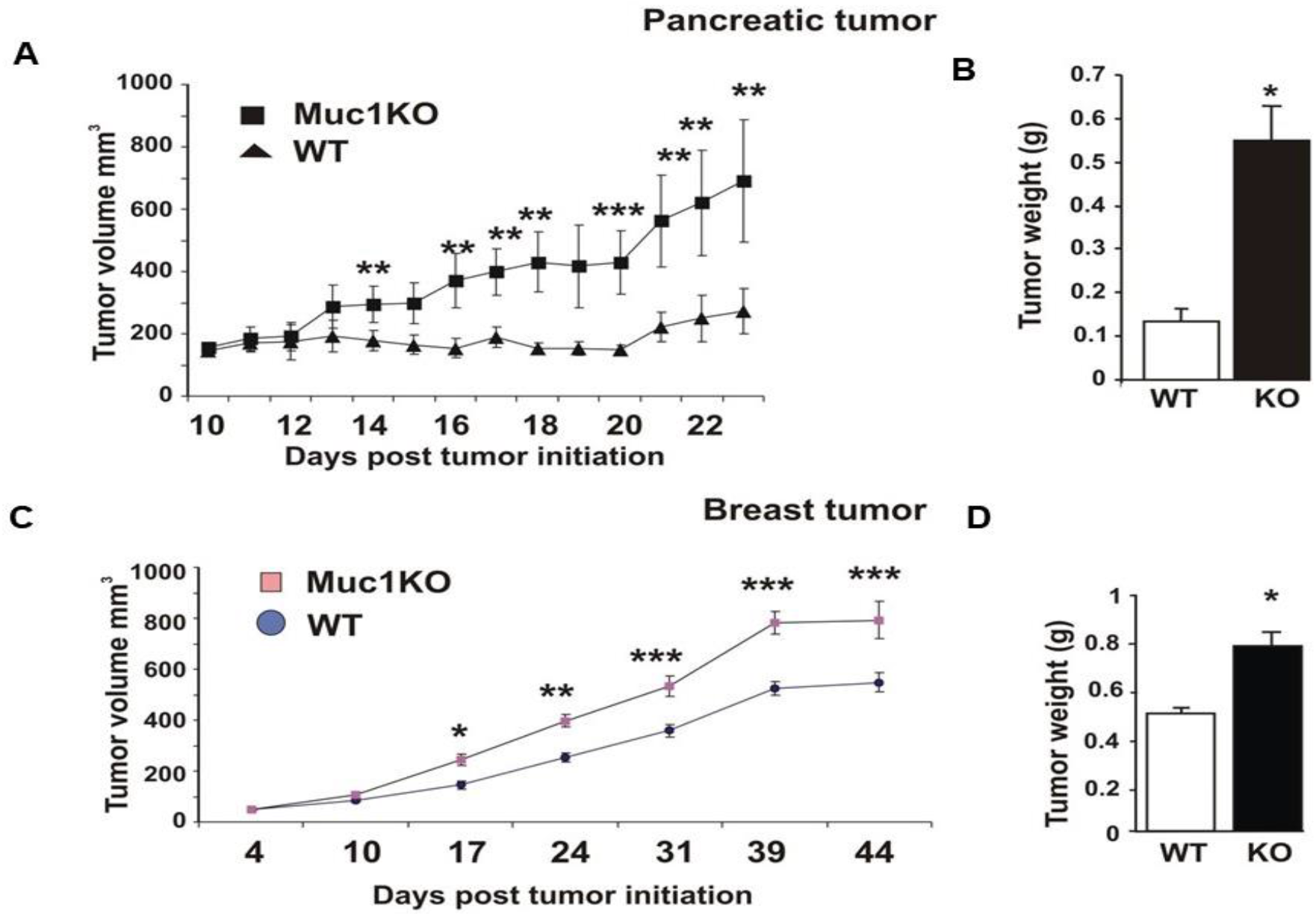
Increased tumor burden in MUC1KO mice. A and C) WT and MUC1KO mice were injected subcutaneously with 10^6^ cells and tumor burden was measured every 2 days. n=8 mice in each WT and MUC1KO mice for Pancreatic Cancer model and n=6 and n=7 mice in WT and MUC1KO groups respectively for Breast Cancer model were used. B and D) Final tumor weight in MUC1KO and WT mice injected with 10^6^ KCKO and C57MG cells. Welch’s t-test was used to compare between WT and MUC1KO groups and p<0.01 was considered to be significant.

### Increased expansion and migration of MDSCs to the spleen of MUC1KO mice and higher levels of TGF-β in their serum compared to that of WT mice

Spleen is the first major lymphatic organ to which MDSCs migrate to exert their suppressive function prior to infiltrating tumors. To study whether MUC1 regulates expansion and migration of MDSCs to the spleen of tumor bearing mice, we analyzed splenocytes from these mice for Gr1+CD11b+ MDSC level by flow cytometry. According to our results, both breast cancer C57MG and pancreatic cancer KCKO cells induced significant expansion/migration of MDSCs into the spleen, and this expansion was significantly higher in the MUC1KO mice versus the WT mice (Figure 2A). Due to increased expansion and suppressive function of MDSCs in MUC1KO mice, and the inflammatory nature of these cells, we wanted to know whether these mice express higher levels of inflammatory cytokine TGF-β in the serum. Serum from tumor-bearing and healthy mice were analyzed for TGF-β secreted by MDSCs which is also known to induce their expansion [45, 46]. Our results show that in both cancer models, levels of TGF-β were significantly higher in MUC1KO mice (Figure 2B, C). Increased suppressive activity of MDSCs is reported to be associated with increased level of CD4+FoxP3+ Tregs cells in tumor environment [21]. We wondered if the lack of MUC1 and its association with increased suppressive function of MDSCs has led to increased levels of other suppressive cells such as T-regs. Indeed, we found that lymphocytes derived from tumor bearing MUC1KO mice contain more CD4+FoxP3+ Tregs cells than those derived from WT mice (Figure 2D). Overall, our results show that during cancer progression, MUC1KO mice have higher levels of the immunosuppressive cytokine TGF-β and increased number of T-regs. This suggests that normal MUC1 on immune cells has a role in influencing the expansion and immunosuppressive function of MDSCs in the tumor microenvironment.

**Fig 2.**
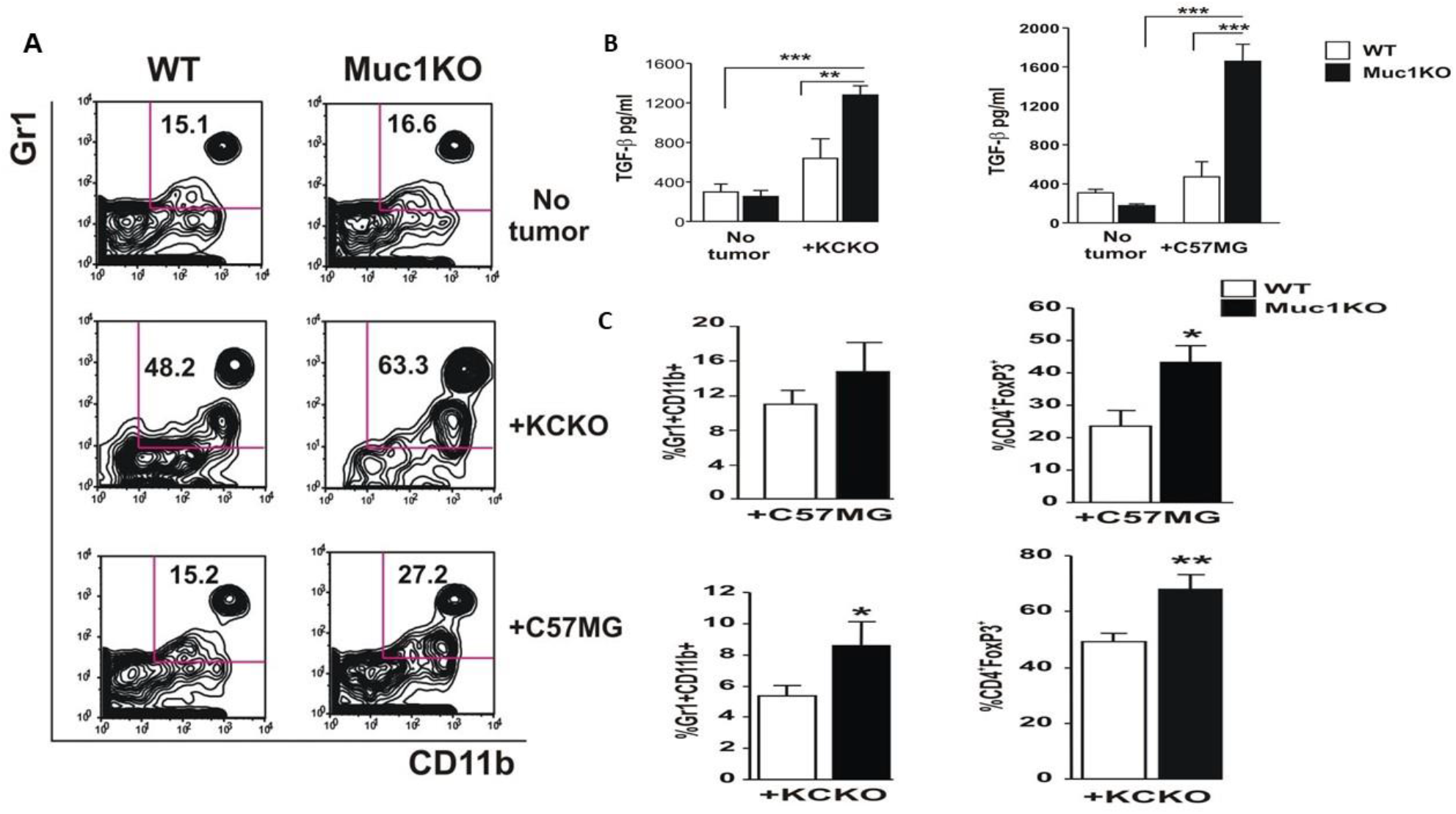
MDSCs from MUC1KO mice have higher frequency and create a more immune suppressive environment. A) Splenocytes from healthy and tumor bearing WT and Muc1KO mice were isolated and labeled with anti Gr1 and CD11b antibodies. Representative plots from three separate experiments are shown. B) Levels of TGF-β was measured in the serum of healthy and cancer bearing WT and MUC1KO mice using ELISA kits. C) Levels of MDSCs and CD4+FoxP3+ cells were measured in the tumor of WT and Muc1KO mice using flow cytometry.

### Spleen-derived MDSCs from MUC1KO mice have higher immunosuppressive phenotype

Phosphorylated STAT3 (pSTAT3) is an enhancer of stemness and mesenchymal properties of MDSCs [47, 48]. We observed an increased level of pSTAT3 in the spleen derived MDSCs from MUC1KO mice compared to WT mice (Figure 3A). Activated STAT3 pathway is known to aid expansion of MDSCs and prevent maturation of myeloid progenitor cells [48]. To further study the mechanism of increased suppressive activity of MDSCs in MUC1KO mice, we wondered whether levels of suppressive enzymes were any different in MDSCs from MUC1KO versus WT mice. Our results showed that levels of ARG1 in the MDSCs of MUC1KO and WT mice bearing the KCKO tumor was increased compared to non-tumor bearing mice (Figure 3E). However, there was no difference between levels of ARG1 between WT and MUC1KO mice bearing the C57MG tumor (Figure 3B). We then hypothesized that even though ARG1 levels in KO and WT MDSCs are not different, it is possible that ARG1 activity is different. Since ARG1 converts L-arginine to urea, we tested the levels of urea production in Gr1+CD11b+ MDSCs sorted from the spleen of WT/KO mice. Surprisingly, we found that urea production in MDSCs from MUC1KO mice was significantly higher than that of its WT counterpart indicating a more suppressive phenotype in MUC1KO mice (Figure 3C).

**Fig. 3.**
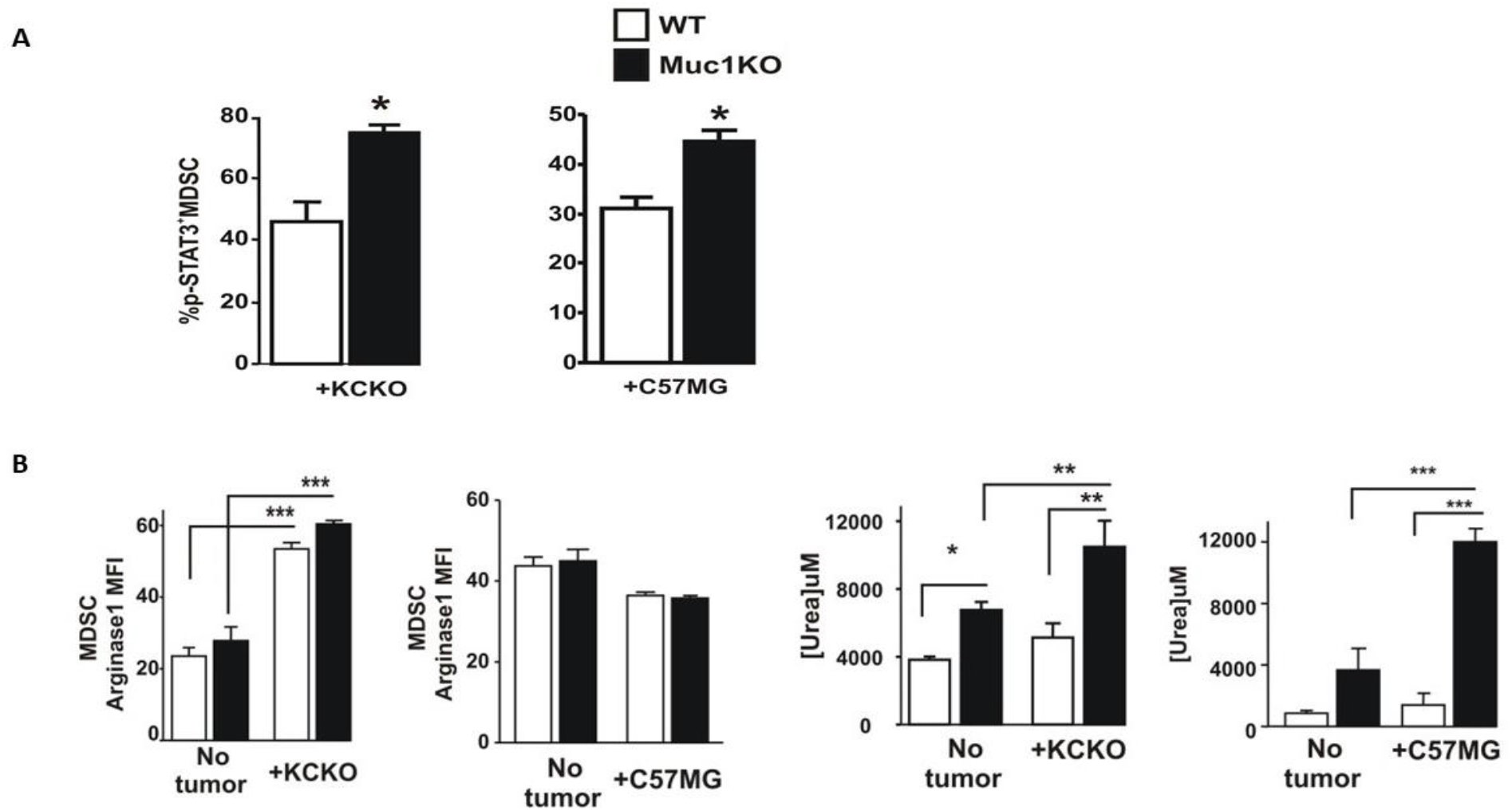
Phenotypic characterization of MDSCs in the BM of healthy and tumor bearing WT and Muc1KO mice. A) Splenocytes from tumor bearing WT and MUC1KO mice were isolated and labeled with antibodies against Gr1, CD11b, and pSTAT3. B) Levels of Arginase-1 expression in MDSCs of WT and MUC1KO mice was measured via flow cytometry. Arginase-1 activity in sorted MDSCs was measured using a urea assay.

### MDSCs from the spleen of MUC1KO tumor bearing mice lead to increased suppression of cytotoxic T-cells compared to WT MDSCs

We hypothesized that differential expression of suppressive enzymes in BM MDSCs might cause MDSCs derived from the spleen of MUC1KO mice to be more effective in T-cell suppression. MDSCs from WT and MUC1KO mice with C57MG and KCKO tumors cells were co-cultured with naïve syngeneic T cells stimulated with α-CD3/CD28 antibodies and supernatants were collected to test for IL-2 and IFN-ɣ production. Our results showed that MDSCs from the spleen of cancer bearing MUC1KO mice were more effective in inducing T-cell suppression indicated by lower IL-2 and IFN-ɣ production by T-cells (Figure 4A-C). Thus, we report that MUC1 regulates the suppressive function of MDSCs under cancer conditions. In agreement with our hypothesis, our data indicate that MUC1 regulates the suppressive functions of MDSCs. Increased ARG1 activity in MUC1KO MDSCs explain the reason behind decreased production of IL-2 and IFN-ɣ by the T-cells observed in Figure 3B.

**Fig. 4.**
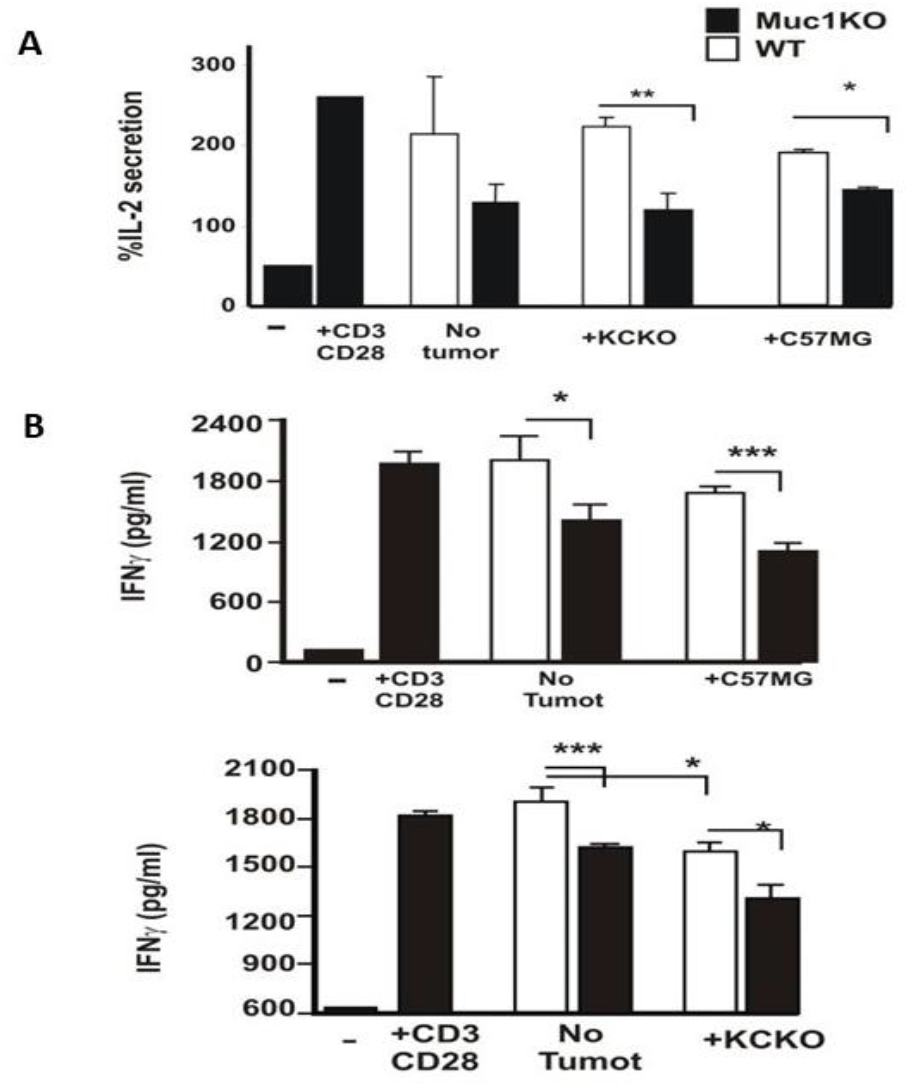
MDSCs from MUC1KO mice inhibit production of proinflammatory cytokines IL-2 and IFN-γ by T-cells. A) MDSCs were sorted from the spleen of WT and MUC1KO mice and incubated with activated T-cells. %IL-2 secretion was measured via ELISA. B) Analysis of IFN-ɣ secretion from CD3/CD28 activated T-cells incubated with WT and MUC1KO MDSCs from healthy and tumor bearing mice.

### BM-MDSCS from MUC1KO mice have different rates of expansion and proliferation compared to that from WT mice

To study whether MUC1 expression on HSCs leads to differential expansion of MDSCs in the bone marrow (BM) of non-tumor bearing mice, freshly isolated BM cells were analyzed for co-expression of Gr1 and CD11b. We found that MUC1KO mice have 15% higher levels of MDSCs in the BM indicating that MUC1 may regulate ontogeny of these cells (Figure 5A).

**Fig. 5.**
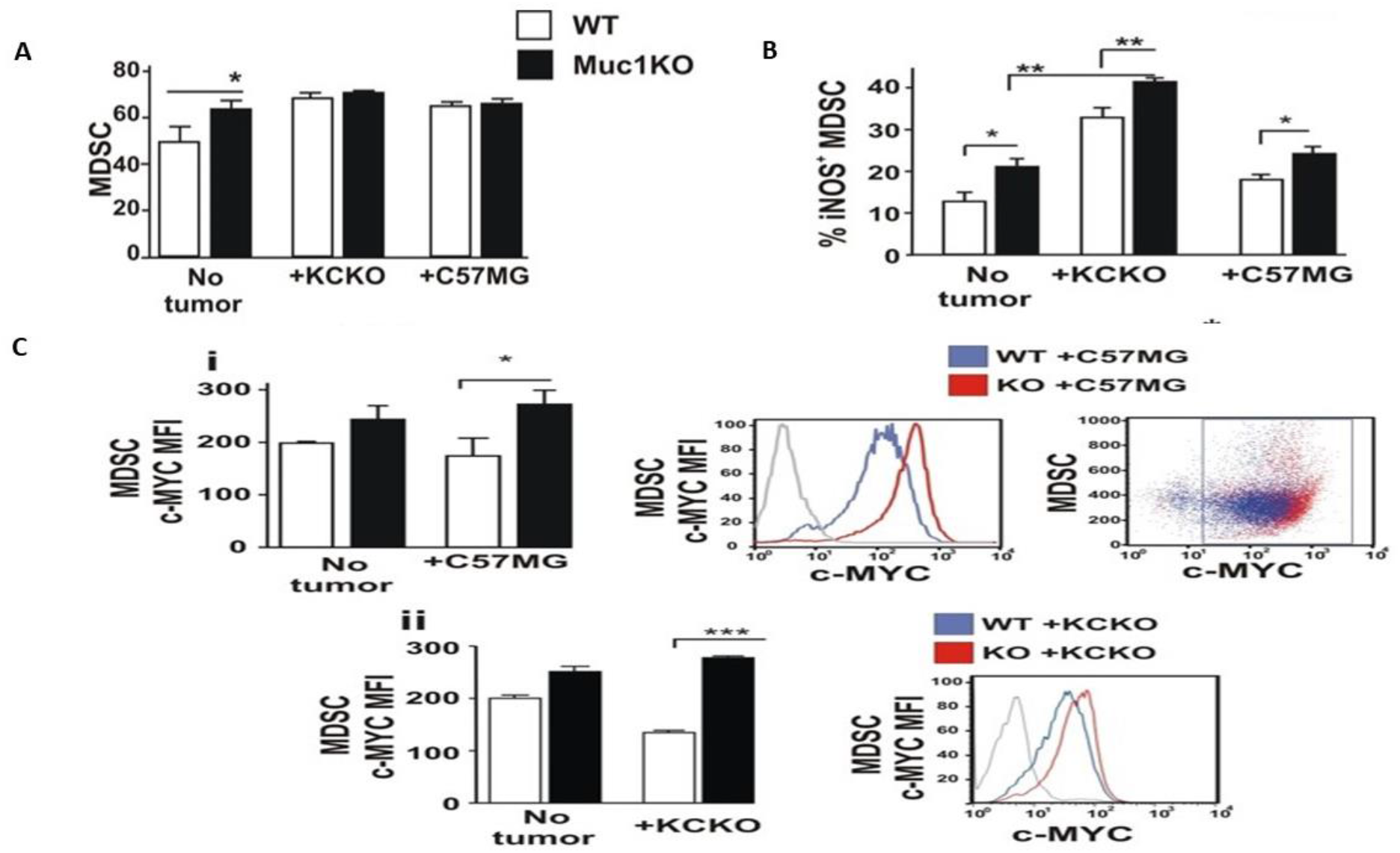
MDSCs from MUC1KO mice produce increased iNOS and c-Myc. A) BM cells from healthy mice were isolated and labeled with antibodies against Gr1 and CD11b. B) Level of iNOS expression by MDSCs from the BM of healthy and tumor bearing mice. C) Level of c-Myc protein expression by MDSCs from the BM of healthy and tumor bearing mice.

**Fig. 6.**
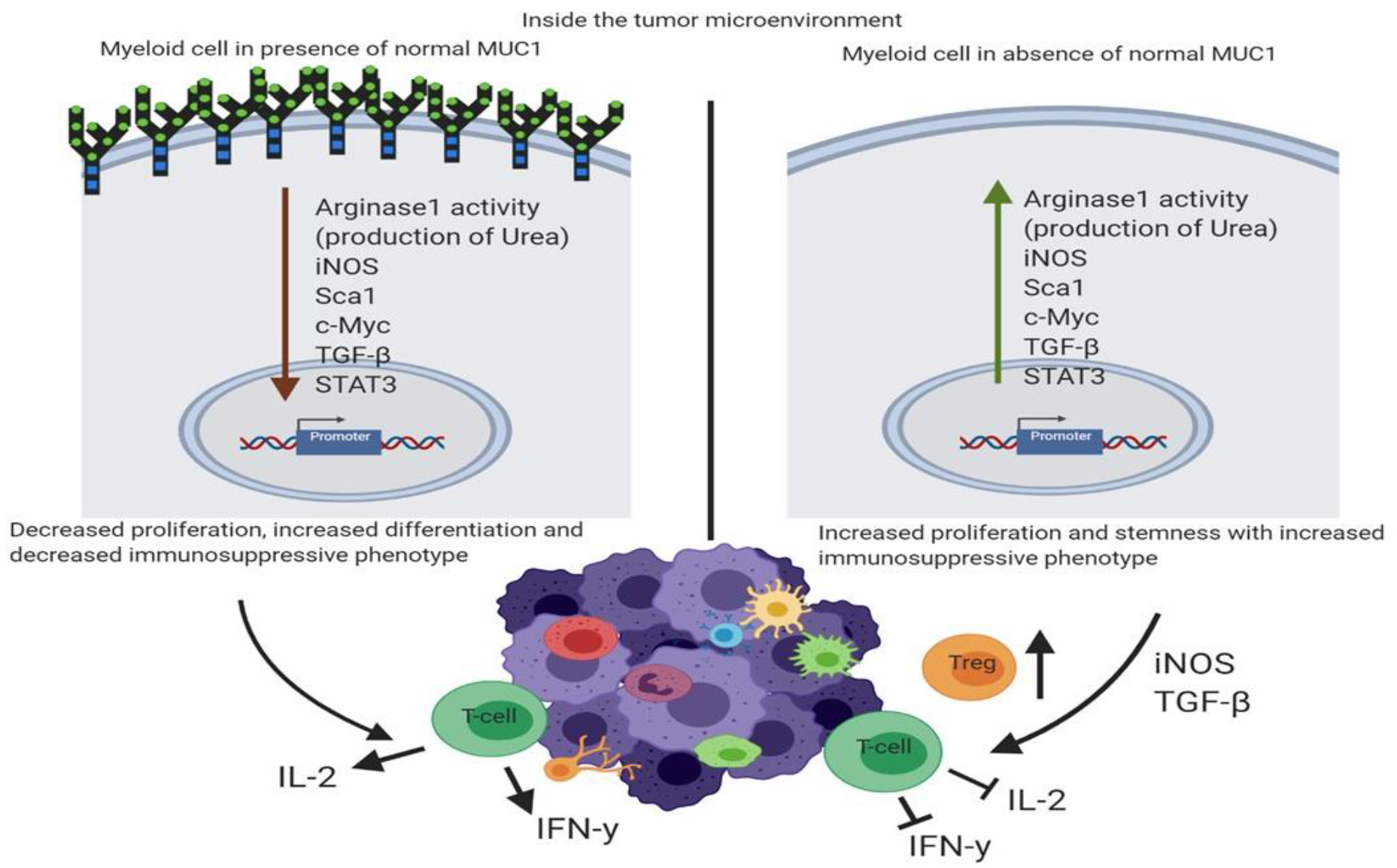
Schematic diagram showing the differences in the tumor microenvironment of a MUC1KO mouse versus that of a WT mouse. Schematic diagram showing the differences in the tumor microenvironment of a MUC1KO mouse (right) vs that of a WT mouse (left). MUC1KO mouse have increased TGF-β and T-reg cells in their tumor microenvironment thus. MDSCs from MUC1KO mouse produce higher Urea, iNOS, stemness marker Sca1, TGF-β and show increased expansion and migration into the tumor, thus inhibiting T-cells from producing IL-2 and IFN-γ.

To characterize the functional activity of BM-MDSCs from WT and MUC1KO mice, we studied levels of iNOS expression by flow cytometry. Our results indicate that higher percentage of MDSCs from healthy non-tumor-bearing MUC1KO mice express iNOS (Figure 5B) than its WT counterpart. Moreover, MDSCs from both breast C57MG and pancreatic KCKO tumor-bearing MUC1KO mice express higher levels of iNOS than MDSCs derived from WT mice suggesting a more suppressive phenotype in MUC1KO MDSCs compared to WT MDSCs (Figure 5B).

To check whether MDSCs from WT and MUC1KO mice differ in their survival and proliferation, we checked the levels of c-Myc protein. c-Myc is responsible for survival and proliferation of MDSCs [49]. We checked levels of c-Myc expression on MDSCs and found that under cancer condition, BM-MDSCs from MUC1KO mice express significantly higher levels of c-Myc than the WT MDSCs (Figure 5C). Our data suggest that in the absence of MUC1 BM-MDSCs are more likely to survive, proliferate, and probably have a longer life span.

Overall, our results have provided evidence supporting the hypothesis that MUC1 regulates the immunosuppressive function of neutrophils (these are not called MDSCs until mice have cancer and in other conditions these are called Neutrophils) in healthy conditions. MUC1 also regulates expansion and suppressive function of Gr^+^CD11b^+^ MDSCs during cancer progression.

## DISCUSSION

Gr1^+^CD11b^+^ MDSC expansion appears in pathologic conditions of chronic inflammation including cancer [23]. It is critical to understand the mechanisms that regulate the functional activity of MDSCs and play a significant role in tumor angiogenesis, drug and immunotherapy resistance, and promotion of tumor metastases. The two groups of signaling previously described to be responsible for accumulation and differentiation of MDSCs include tumor –derived growth factors (STAT3, IRF8, C/EBPβ, Notch, adenosine receptors A2b signaling, NLRP3, Rb1) and proinflammatory cytokines produced by tumor stroma (NF – κβ pathway, STAT1, STAT6, PGE2, COX2) [22]. At the same time, there are fewer reports about downstream targets of tumor derived factors such as MUC1 that contains binding sites for NF – κβ in its promoter and that depends on STAT factors for its expression (reviewed in [41, 50]). The latter finding defines that development of BM progenitors into MDSCs was mediated through down-regulation of β-catenin levels in the absence of MUC1 *in vitro* [30]. Here, we report that MUC1 regulates proliferation and suppressive phenotype of MDSCs in both healthy and cancer conditions. Using tumor models, we were able to show how MDSCs affect tumor microenvironment and immune response under MUC1 null condition.

We have shown high level of MUC1 expression in normal hematopoietic stem cell and progenitor compartment (Supplementary Figure 1) confirming previously published data [51].

Previous studies showed that MDSCs influence differentiation of CD4+ cells to Treg cells through the production of TGF-β cytokine [52, 53]. Here, we report an increased level of TGF-β secretion by the MDSCs in MUC1KO condition with enrichment of Treg cells in tumor infiltrating lymphocyte population. Observed MUC1-dependent Treg proliferation and suppression of T-cell function are responsible for the enhanced tumor growth in both the pancreatic KCKO and breast C57MG cancer models *in vivo*.

We also found that MUC1 knockdown condition was associated with higher proliferation of MDSCs and more immunosuppressive phenotype. These effects may be mediated through the detected increased levels of pSTAT3 and c–Myc that are responsible for the survival, immature phenotype and immunosuppressive features of MDSCs [22, 48, 54].

Future research will be focused on understanding the mechanisms of MUC1-regulated expression of c-Myc in MDSCs. It was reported that MUC1 may regulate c-Myc expression on transcriptional level [55, 56]. Previously, it was reported that MUC1 C-terminal subunit (MUC1-C) activated the WNT/β-catenin pathway and promoted occupancy of MUC1-C/β-catenin/TCF4 complexes on the c-Myc promoter in non-small cell lung cancer [55]. A similar mechanism for activation of c-Myc by MUC1-C/β-catenin/TCF4 complexes was detected in multiple myeloma [56]. Thus, MUC1 may affect c-Myc expression on the transcriptional level. Alternative mechanisms of MUC1 regulation includes STAT3-mediated overexpression of c-Myc. STAT3 is known to control the granulocyte colony-stimulating factor (G-CSF) - mediated induction of CCAAT-enhancer-binding protein beta (C/EBPβ), a crucial factor in the emergency granulopoiesis response, in myeloid progenitor cells [57]. For instance, G-CSF driven STAT3 activation resulted in c-Myc overexpression by increasing the binding of C/EBPβ to c-Myc promoter [57]. Thus, STAT3 can induce expansion of MDSCs via upregulation of C/EBPβ [22]. Here, we report increased levels of pSTAT3 in the spleen-derived MDSCs from MUC1KO tumor-bearing mice compared to those from WT tumor-bearing mice. MUC1-mediated increased activity of pSTAT3 deregulates expression of c-Myc and may be a reason behind the observed overexpression of c-Myc in MDSCs. Further research is needed to study the effect of knocking down MUC1 on downstream targets in c-Myc target genes, such as telomerase reverse transcriptase, cyclin-dependent kinase 4, Cyclin D2 and glutamate—cysteine ligase catalytic subunit [55]. Currently, the most described effects of MUC1 regulation of c-Myc and STAT3 were observed using cancer cells models [41]. In breast cancer cells, the MUC1-C was found to be necessary for gp130-Janus-activated kinase 1 (JAK1)-mediated STAT3 activation [58]. However, an open question remains about the specific role of MUC1 in cancer cells versus in MDSCs in regulating target genes and the role of posttranslational modification of mucins in this process.

In this study, for the first time we report the critical role of MUC1 in regulating the suppressive phenotype of MDSCs in tumor microenvironment. Lack of MUC1 in MDSCs correlated with increased levels of iNOS, ARG1 activity and TGF-β secretion that are markers of MDSCs immunosuppressive activity [59]. It is reported that iNOS induces NO leading to inhibition of T-cells via downregulation of IL-2 production and receptor signaling and induction of T-cell apoptosis [60]. ARG1 - dependent depletion of L-arginine impair T-cell proliferation and function [24, 52, 60]. We were able to register significant impact of MUC1-mediated increasing iNOS and ARG1 activity on T-cell function including repression of IL-2 and IFN-ɣ production. Thus, knockdown of MUC1 in MDSCs may have a negative effect on the immune response against the tumor [8].

In summary, MUC1 plays a critical role in regulating the immunosuppressive functions of MDSCs in the tumor microenvironment, thus, in turn affecting carcinogenesis. However, some critical questions still need to be answered regarding MUC1 expressing MDSCs and their role in tumor progression and metastasis. It was demonstrated that STAT3 activation in MDSCs is required to increase the cancer stem cell population in pancreatic adenocarcinoma [31]. CSCs (Cancer Stem Cells) are resistant to conventional treatments and are considered the reason behind cancer metastases and recurrence after clinical remission [61]. Further studies are required to assess the effect of knocking down MUC1 in MDSCs on cancer stemness in a STAT3– dependent manner.

Previously, the paradoxical roles of MUC1 based on its pattern of glycosylation in normal versus cancer cells have been reported in case of cancer-associated infections [62]. Our data show that in future, specific targeting of tumor-associated MUC1 (tMUC1) without targeting the normal MUC1 on immune cells should be a preferred strategy to increase efficacy against tumors. Here we report that normal MUC1 on the MDSC populations reduce their expansion, proliferation, migration to the spleen and immunosuppressive activities. This should not be confused with the tumor-associated MUC1 (tMUC1) which when expressed on tumor cells increases their aggressive phenotype and is a marker of poor prognosis. We are the first to report that nonspecific elimination of MUC1 in MDSCs leads to immature immunosuppressive phenotype of MDSCs and potentiates tumor growth. On these terms, a recently reported novel monoclonal antibody TAB004 (OncoTAb, Inc., Charlotte, NC), which specifically targets the hypoglycosylated/tumor-associated form of MUC1 (tMUC1) seems to be a perfect delivery agent for targeting tMUC1 with drugs and peptides [37, 63, 64].

## Author Contributions

Data curation, investigation and analysis: MS, LR, SN and MB; data analysis, preparation of manuscript and revision: MS, LR, MB and CD; supervision, funding and revision: PM.

## Acknowledgments

We thank Dr. Chandra Williams, DVM, DACLAM, CPIA, University Veterinarian and Vivarium Director and all vivarium staff members.

## Supplemental Materials

**Supplemental Fig 1.**
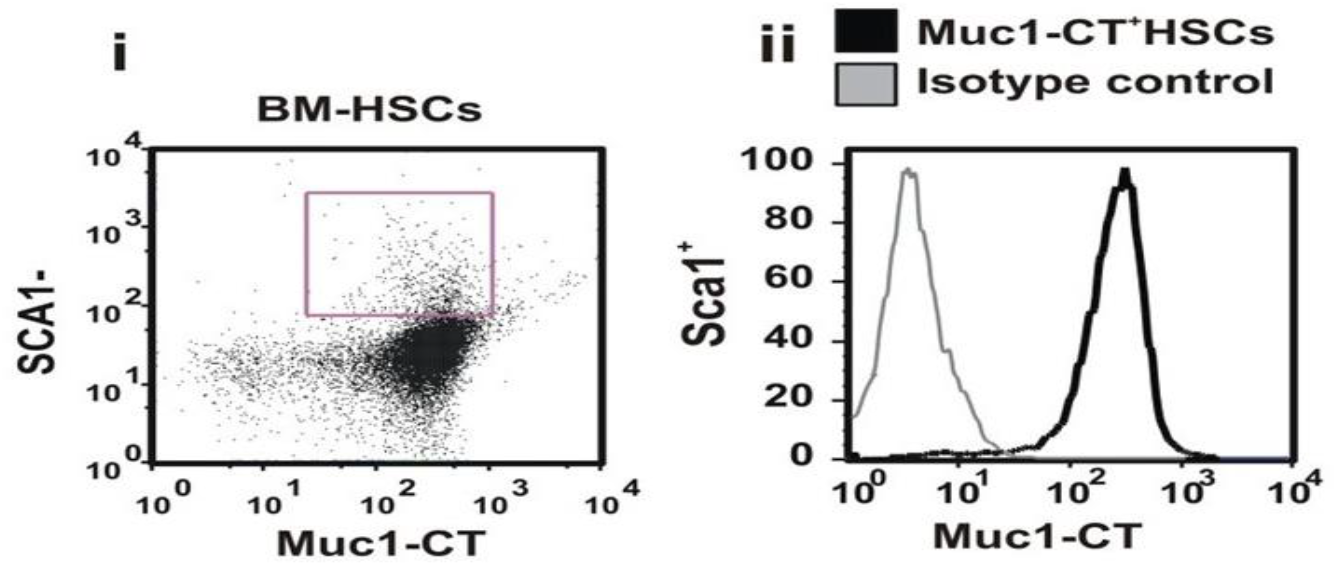
HSCs and MDSCs express high levels of MUC1. BM cells are co-stained with antibodies directed against Sca1 and Muc1-CT. Data are shown as both percentages (i) of Sca1+ cells expressing Muc1-CT and (ii) the expression level of this protein by these cells.

