## Supplementary figures and images for "Repression of MUC1 promotes expansion and suppressive function of myeloid-derived suppressor cells in pancreatic ductal adenocarcinoma and breast cancer murine models"

### Supplemental Figure 1

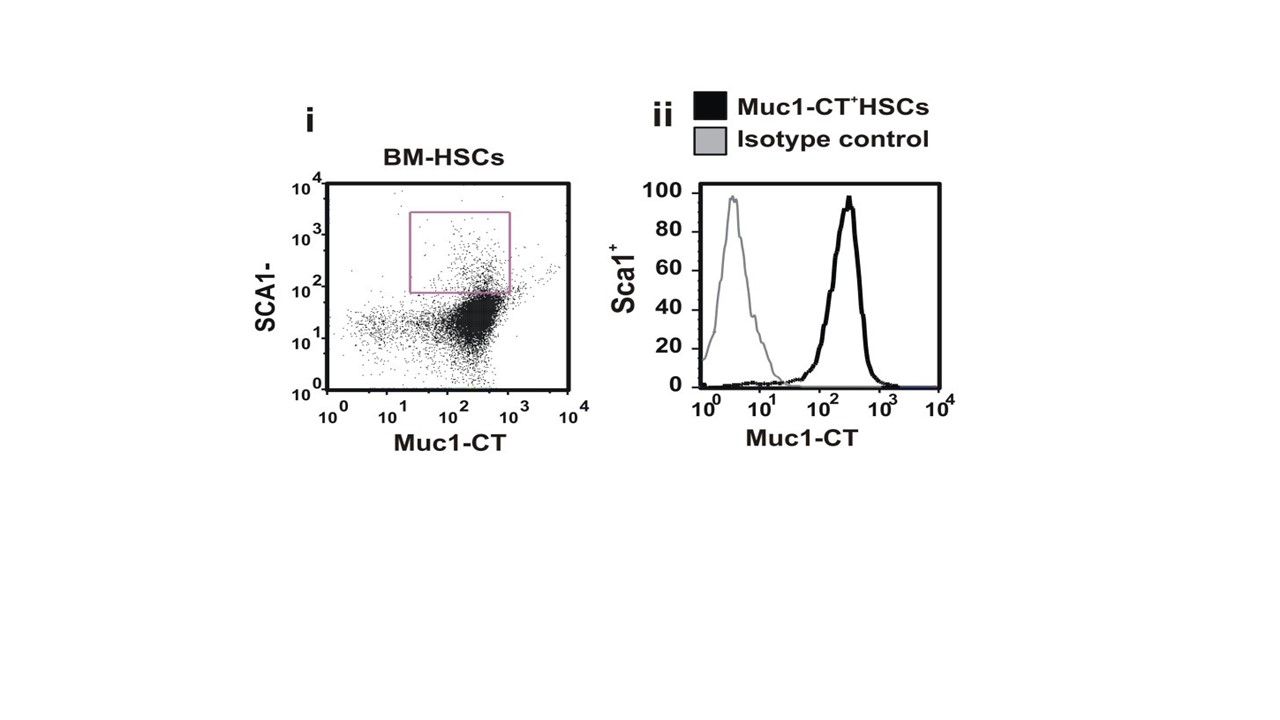
